# The effect of a short oxygen exposure period on algal biomass degradation and methane formation in eutrophic and oligotrophic lake sediments

**DOI:** 10.1101/2024.04.04.588097

**Authors:** Sigrid van Grinsven, Natsumi Maeda, Clemens Glombitza, Mark A. Lever, Carsten J. Schubert

## Abstract

Eutrophication is suggested to enhance lacustrine methane emissions, due to enhanced sedimentary decomposition rates of algal biomass, and more frequent occurrence of water column anoxia. We investigated methane emissions from sediments originating from both a eutrophic and oligotrophic lake, and tested the effect of additional algal C inputs. Additionally, we investigated the effect of a pulse supply of oxygen, a mediating measure that is currently being used in the investigated eutrophic lake. Our results show a large legacy effect of eutrophication, but the methane release from new algal biomass additions was the same, although the process proceeded more rapidly in the eutrophic sediments. A short, 3-week pulse of oxygen lowered the emitted methane from both types of sediments by 50%, not only reducing the emissions of algal biomass additions, but also reducing methane emissions from the experiments without fresh organic matter inputs. This effect was relatively long-lasting: its effects were visible for several weeks after anoxic conditions were re-established, making it a potentially interesting measure to lower methane emissions over a longer period. Volatile fatty acid concentrations in the sediments were lowered due to oxygen exposure. Both the methanogenic and methanotrophic community composition showed surprisingly little response to the oxygen or algal biomass pulses. Overall, our results show that providing sediments with a brief pulse of oxygen following an algal bloom event, could strongly mediate the methane emissions following such an event. Such measures could be considered by policy makers to limit greenhouse gas emissions from managed lakes.

## Introduction

Lakes are known to be significant contributors to global methane emissions, despite their relatively small surface area (Bastviken et al. 2004). Methane emissions are the result of the net outcome of two processes: the production of methane, called methanogenesis, and the consumption of methane, methanotrophy. Methane emissions from aquatic environments originate mostly from sediments. Organic matter is delivered from either internal lacustrine (autochthonous) or external, e.g. riverine and terrestrial (allochthonous) sources. Autochthonous primary production, e.g. by planktonic microalgae in the water column, captures CO_2_ from the atmosphere to produce biomass. After cell death, this biomass sinks down the water column, and becomes part of the sediment. The decomposition of this biomass lowers dissolved oxygen concentrations in bottom water and surface sediments and frequently enhances sedimentary methane production (Fiskal et al., 2019; van Grinsven et al., 2022). Eutrophication, the increase in (mainly algal) primary production due to increased nutrient concentrations in lakes, has thus been shown to increase methane emissions from lakes (Beaulieu et al. 2019).

Most sublittoral lake sediments are anoxic from a depth of a few mm to cm below the sediment surface. This is due to the mainly diffusive transport of oxygen into sediment and the high rates of aerobic decomposition processes at the sediment surface. In the underlying anoxic sediment, organic matter breakdown is performed by a community of hydrolytic, fermentative, and respiring microorganisms. Generally, methanogenesis is expected to occur only after the depletion of other, more energy-rich anaerobic oxidants, such as nitrate, nitrite, metal-oxides and sulfate (Bastviken et al. 2004), though strong overlaps in the distribution of methanogenesis with other anaerobic respiration reactions have also been observed (Fiskal et al., 2019).

Generally, the anaerobic breakdown of organic matter follows three steps: first, the large, complex organic compounds (e.g. macromolecules, polymers) are broken down to their building blocks (e.g. monomers, oligomers, fatty acids) by extracellular reactions (e.g. hydrolysis). Subsequently, these building blocks are taken up by microbial cells and fermented to smaller chemical compounds, such as H_2_, alcohols (e.g. methanol, ethanol) and volatile fatty acids (VFAs; e.g., acetate, propionate, butyrate, isovalerate, formate, and pyruvate). These smaller compounds can then undergo a secondary fermentation step by syntrophic microorganisms, prior to respiration, or be directly respired to CO2 or methane using nitrate, nitrite, metal-oxides, sulfate, or CO2 as electron acceptors. Methanogens, which are respiring organisms that gain energy through the production of methane, are mostly obligate anaerobes (Whitman et al. 2014), although their tolerance to oxygen is debated and may differ between clades (Kiener and Leisinger 1983; Zinder 1993; Kato et al. 1993). Methanogenesis in sediments is believed to proceed mainly via three different pathways: CO2 reduction using H2 or formate as electron sources, acetoclastic involving the disproportionation of acetate into CO2 and methane, or methylotrophic methanogenesis, which involves the conversion of methylated compounds, such as methanol, methylamines and methyl sulfides to methane. With few exceptions, these pathways are performed by distinct taxa of methanogens.

A significant fraction of methane is consumed by methanotrophy, the process of methane consumption. Methanotrophy can occur both aerobically or anaerobically via the reduction of various anaerobic electron acceptors. Methanotrophs are found both within the archaeal and bacterial domain and include strict aerobes, facultative anaerobes, and obligate anaerobes. Methanotrophic activity often peaks at the oxic-anoxic interface in either the sediment or water column, presumably due to the high energy yields of aerobic methanotrophy.

Past research indicates that high organic matter inputs to lake sediments following algal blooms result in increases in methane concentrations in the sediment (Schulz and Conrad 1994). Both eutrophication and reduced water column mixing due to increased thermal stratification as a result of global warming will likely increase the frequency of algal blooms and contribute to more widespread bottom water anoxia in the future (Hou et al. 2022). While this promotes the deposition and burial of organic carbon in lake sediments, it will also increase methane emissions by increasing methanogenesis rates and lowering rates of aerobic methanotrophy. In addition to future algal blooms, legacy effects, e.g. continued high rates of methane production sustained by the decomposition of older organic carbon from past periods of eutrophication may contribute to these elevated methane emissions. These increases in methane emissions may be a lesser concern for oligotrophic sediments, which are generally lower in organic carbon content and hence methane production rates.

In order to mediate the effects of current and past eutrophication, artificial aeration is applied to lakes in Switzerland. This aeration reduces the detrimental ecological and socioeconomic consequences of seasonal anoxia. In addition, artificial aeration may lower methane emissions from lakes by promoting aerobic methanotrophy and reducing methane production in deep water columns and surface sediments, though the efficacy of artificial aeration in achieving lower methane emissions is not known. Here, we experimentally test the impact of artificial aeration on methane emissions under different trophic regimes by short, pulse-wise, oxygen supply to sediments from oligotrophic Lake Lucerne and eutrophic Lake Baldegg (both Switzerland). Based on slurry and whole-core incubations, we study the impact of oxygen pulses on methane emissions from sediments with and without an initial spike of algal biomass to mimic the situation shortly after an algal bloom. Samples were extracted for gas concentration and isotope analysis, as well as VFA and microbial community analyses. Our results show that oxygen exposure had effects lasting past the oxic period such as that methane emission rates remained lower for up to 10 weeks. The methanogenic and methanotrophic communities did not seem affected by the oxygen exposure.

## Methods

### Study sites

Lake Lucerne is located at the northern alpine front in Central Switzerland (47°N, 8°E, 434 m a.s.l). It has a surface area of 116 km^2^, and is fed by four alpine rivers that provide ±80% of the lakes total water supply (Schnellmann et al. 2002). It is oligotrophic, with a maximum P-concentration within the past century of 1.7 μM (Bürgi and Stadelmann 2002).

Lake Baldegg is located in an area with intensive cattle and pig farming within Central Switzerland. It has a surface area of 5.22 km^2^. Eutrophication has been ongoing, with a peak in the 1970s, reaching P concentrations of 15.4 μM before remediation measures were put in place. It has had an anoxic hypolimnion for almost 100 years until artificial aeration was started in 1982 (Gächter and Wehrli 1998). A map showing the location of both lakes is shown in Fig. S1.

### Field sampling

Lake Lucerne sediment cores were taken in November 2020 and February 2021 at, 47.00085N, 8.33697E, water depth ± 70 m. Lake Baldegg sediment cores were taken in June 2021 from the center of the lake (47.193071N, 8.265238E). Both lakes were sampled with a multicorer device containing 10 cm diameter, transparent butyrate plastic core liners (liner length?). The average core length was 40 cm. All sediment cores were brought into a climate room of 10°C within 2 hours after core collection, and stored until further processing.

### Slurry incubation experiments

An overview of the experiments is presented in Table 1. All slurry experiments were performed in triplicate. To set up the slurry experiments, bottom water was taken from the cores prior to slicing, to be used for the slurries. For the oxic incubation experiments, the top 5 cm of one core was divided into three parts (43 ml) and disposed into 0.5L Schott bottles that each contained 43 ml of bottom water, corresponding to a sediment:water ratio of 1:1. Each bottle was closed with an adapted stopper, containing a three-way-stopcock that could be opened to allow throughflow of sediment slurry and gas without exchange with the air.

**Table 1.**
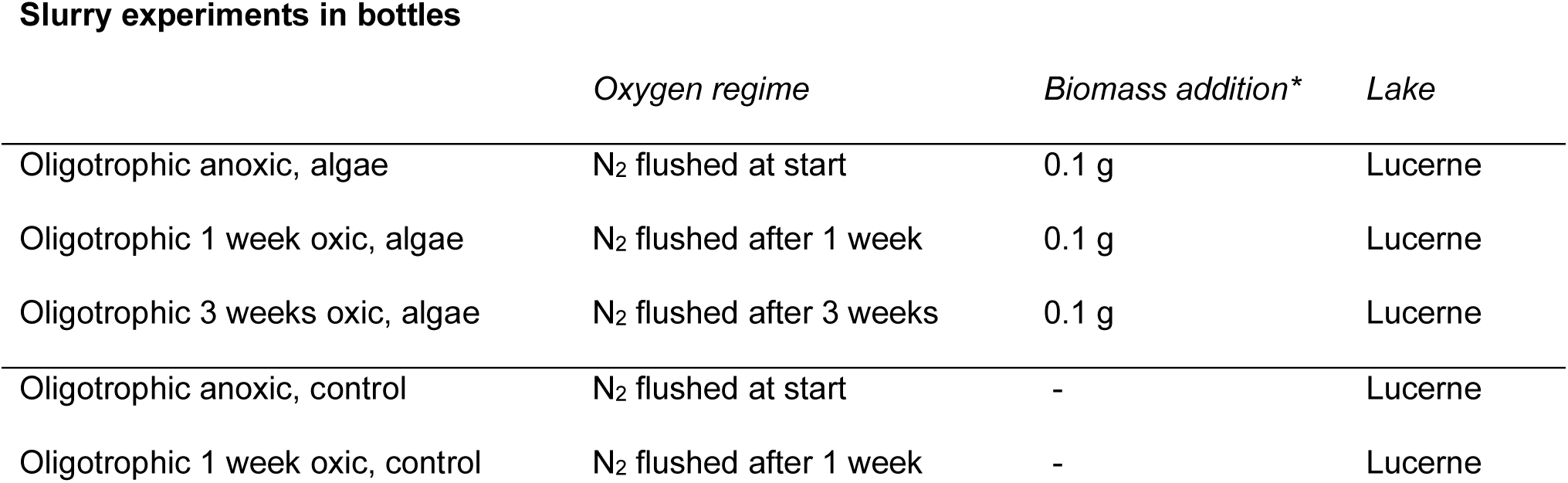
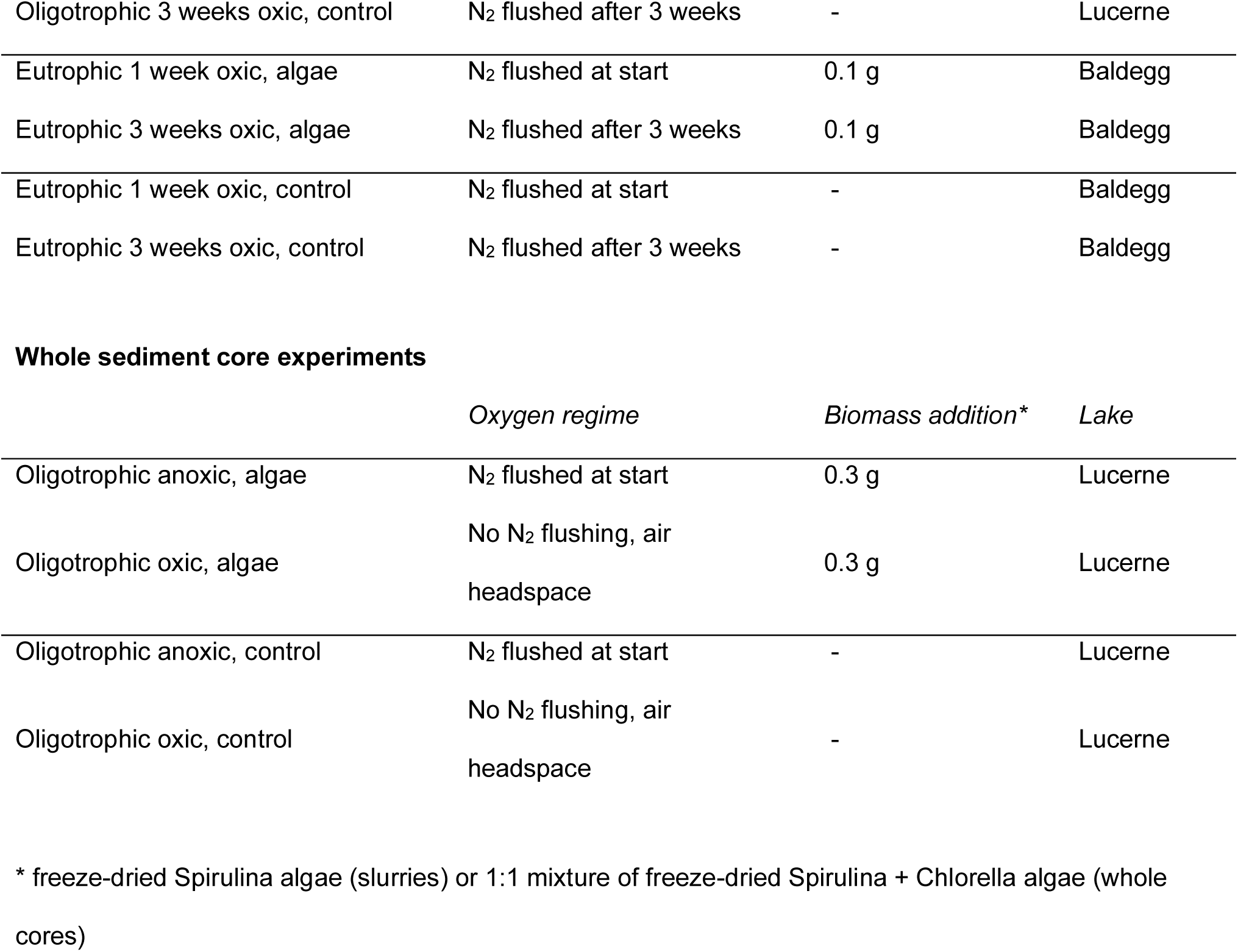
Overview of experiments. Anoxic slurry incubations directly contained sediment from 0 – 15 cm depth, whereas initially oxic slurries only received 0 – 5 cm material initially, and received the anoxic 5 – 15 cm material at the timepoint of N_2_ flushing, as is described in the Methods section.

**Slurry experiments in bottles**
|  | *Oxygen regime* | *Biomass addition\** | *Lake* |
| --- | --- | --- | --- |
| Oligotrophic anoxic, algae | N<sub>2</sub> flushed at start | 0.1 g | Lucerne |
| Oligotrophic 1 week oxic, algae | N<sub>2</sub> flushed after 1 week | 0.1 g | Lucerne |
| Oligotrophic 3 weeks oxic, algae | N<sub>2</sub> flushed after 3 weeks | 0.1 g | Lucerne |
| Oligotrophic anoxic, control | N<sub>2</sub> flushed at start | - | Lucerne |
| Oligotrophic 1 week oxic, control | N<sub>2</sub> flushed after 1 week | - | Lucerne |
| Oligotrophic 3 weeks oxic, control | N<sub>2</sub> flushed after 3 weeks | - | Lucerne |
| Eutrophic 1 week oxic, algae | N<sub>2</sub> flushed at start | 0.1 g | Baldegg |
| Eutrophic 3 weeks oxic, algae | N<sub>2</sub> flushed after 3 weeks | 0.1 g | Baldegg |
| Eutrophic 1 week oxic, control | N<sub>2</sub> flushed at start | - | Baldegg |
| Eutrophic 3 weeks oxic, control | N<sub>2</sub> flushed after 3 weeks | - | Baldegg |

Table 1. Overview of experiments.
|  | *Oxygen regime* | *Biomass addition\** | *Lake* |
| --- | --- | --- | --- |
| Oligotrophic anoxic, algae | N<sub>2</sub> flushed at start | 0.3 g | Lucerne |
| Oligotrophic oxic, algae | No N<sub>2</sub> flushing, air headspace | 0.3 g | Lucerne |
| Oligotrophic anoxic, control | N<sub>2</sub> flushed at start | - | Lucerne |
| Oligotrophic oxic, control | No N<sub>2</sub> flushing, air headspace | - | Lucerne |

\* freeze-dried Spirulina algae (slurries) or 1:1 mixture of freeze-dried Spirulina + Chlorella algae (whole cores)

For the anoxic bottles, slurries were prepared in 1L bottles, containing 400 ml N_2_-flushed anoxic bottom water, to which the upper 15 cm of one core was added. The contents were mixed by shaking, again flushed with N_2_ to reduce the amount of oxygen, and then distributed anoxically over 3 N_2_ flushed 0.5L Schott bottles. Each bottle was closed with the same adapted stoppers described for the oxic experiments. The procedure was repeated once, resulting in 6 anoxic sediment slurry bottles. t_0_ samples (samples taken at the beginning of the experiment) were taken from the second sediment slurry and were immediately frozen at −20°C. Freeze-dried algal biomass (0.1 g of freeze-dried Chlorella; DietFoods CH) was added to a subset of the bottles. Incubations proceeded at 10°C in the dark. The oxygen concentration was measured daily to bi-weekly in 8 out of the 12 oxic bottles, using optical oxygen sensor spots (Pyroscience, UK).

After 1 week or 3 weeks, the oxic bottles were flushed with N_2_ for 15 minutes to remove oxygen, after which they received additional anoxic sediments, as explained below. The resulting oxygen trends are shown in Fig. S2. For each 3 bottles, one 1L bottle with slurry was prepared: sediments of the 5 – 15 cm depth interval of one, previously unsliced, core were mixed with 270 mL bottom water. Each incubation bottle received 1/3 of the contents, making the total sediment content of the 1-week oxic, 3-week oxic and anoxic bottles equal. Sediments were added via the sampling ports in the stoppers.

At each gas sampling time point, headspace gas was sampled via the sampling port in the bottle caps. Prior to gas extraction, 10 ml of N_2_ or air was pushed into the bottle, to clean the sampling port. Immediately after, 10 ml of headspace gas was sampled and stored in N2 flushed 70 ml serum bottles. At each VFA (volatile fatty acids) and DNA sampling time point, 1.5 ml overlying water and 1.5 ml mixed slurry were sampled as described below. For the VFA samples, samples without particulates were required.

To achieve this, the bottles were carefully tipped over, so the natural layering of sediment at the bottom and water at the top remained. The water was then sampled via the sampling port. For the DNA sample, the bottle was briefly shaken and then held upside down to take the mixed water + sediment DNA sample. Both acetate and DNA samples were put directly on ice, and after the sampling series was finished, moved to −20°C (acetate) or −80°C (DNA) freezers.

### Whole core incubation experiments

Whole core experiments were performed in quadruplicate. To prepare each core for incubation, a 12 cm headspace was created, leaving approximately 10 cm bottom water above the sediment-water interface of each core. One core of each treatment received an oxygen sensor spot (Pyroscience, UK), glued to the inner wall of the core liner. Ca. 18 hours after sampling, freeze-dried algal biomass was added to selected cores (0.3 g per core, 1:1 mixture of freeze-dried Chlorella and Spirulina; DietFoods, CH), corresponding to the same amount of algal biomass per sediment mass as in the slurry incubations, and following (Dai et al. 2005; Hiltunen et al. 2021). The algal biomass was provided to the sediments by careful deposition on the sediment surface using a pipet, without disturbing the sediment-water interface. Adjusted rubber stoppers with sampling ports were used to seal the cores. The headspace and overlying water of the cores for the anoxic incubations were flushed for at least 10 minutes with N_2_. All cores were then placed at 10°C in the dark. Oxygen concentrations were measured at the start of the experiment and at irregular intervals over the course of the experiment, shown in Fig. S3. Over the course of the experiment, gas samples for CH_4_, CO_2_ and N_2_O analysis were taken via the sampling ports. 10 ml of the gas headspace was extracted and placed into 70 ml N_2_-flushed serum bottles, closed off with butyl stoppers. After the gas sampling, 10-15 ml of N_2_ gas (anoxic cores) or N_2_ or air (oxic cores) was added via the sampling ports, to equilibrate the internal and external pressure and limit the risk of leakage or contamination. The gas pressure prior to sampling was determined with a pressure meter and noted for each timepoint (not shown).

### Gas and VFA analysis

Gas samples were analyzed by gas chromatography (GC; Agilent 6890N, Agilent Technologies) using a Carboxen 1010 column (30 m x 0.53 mm, Supelco), a flame ionization detector and an auto-sampler (Valco Instruments Co. Inc.).

Volatile fatty acids (VFAs) were filtered through pre-cleaned syringe filters Acrodisc™, 0.2 µm PES membrane, Supor™) and analyzed by two dimensional ion chromatography (2D IC) at ETHZ according to the method described in (Glombitza et al. 2014) with some modifications. The instrument used was a Dinonex™ ICS6000 (Thermo Fisher Scientific) equipped with two 2.5-mm columns (AS24 for the first dimension and AC11HC for the second dimension). The Retention time window on the first IC dimension to collect the bulk VFAs for injection onto the second IC column was set to 3 min – 6.5 min to account for the low salinity of the freshwater samples compared to the original method, as described in (Schaedler et al. 2018; Vuillemin et al. 2023). Likewise, the VFA standards for quantification (mixed standards of formate, acetate, propionate, butyrate, valerate, isovalerate and pyruvate at 1, 5, 10, 50 and 100 µmol L^-1^) were prepared in Milli-Q^®^ water instead of IAPSO seawater as described in the original method. Quantification was done using the conductivity detector signal of the second IC dimension.

### Microbial community analysis

Each DNA sampled was stored at −80°C until processing. DNA was extracted using the Qiagen Powersoil DNeasy kit without adaptations. The DNA concentration of all extracts was measured on a Nanodrop device (Thermo Scientific). When the concentration was below 2 ng/μl, an additional extraction was performed, and sampled were pooled. DNA extracts from all experiments were combined in two lanes, including extraction blanks, and send for 16S rRNA NovaSeq PE250 sequencing (30K tags per sample) to Novogene UK, using the general 16S rRNA archaeal and bacteria primer pair 515F and 806R, targeting the V4 region (Caporaso et al. 2012). Quality control and species annotation were performed using the standard Novogene pipelines (https://www.novogene.com/eu-en/services/research-services/metagenome-sequencing/16s-18s-its-amplicon-metagenomic-sequencing/). Raw sequencing data is deposited in the public repository [to ADD].

## Results

Incubation experiments were set up with sediments from oligotrophic Lake Lucerne and eutrophic Lake Baldegg. An overview is presented in Table 1. All slurry experiments emitted significant quantities of methane, except for the temporarily, 3-week oxic (from here on, called ‘oxic’) oligotrophic control slurries. All given concentrations and concentration increases are given in μM per liter headspace volume. The anoxic background methane emission in the eutrophic control slurries exceeded those of the oligotrophic control slurries over 12-fold (85 versus 6.9 μM per week, respectively, Fig. 1; Table S1).

**Fig. 1.**
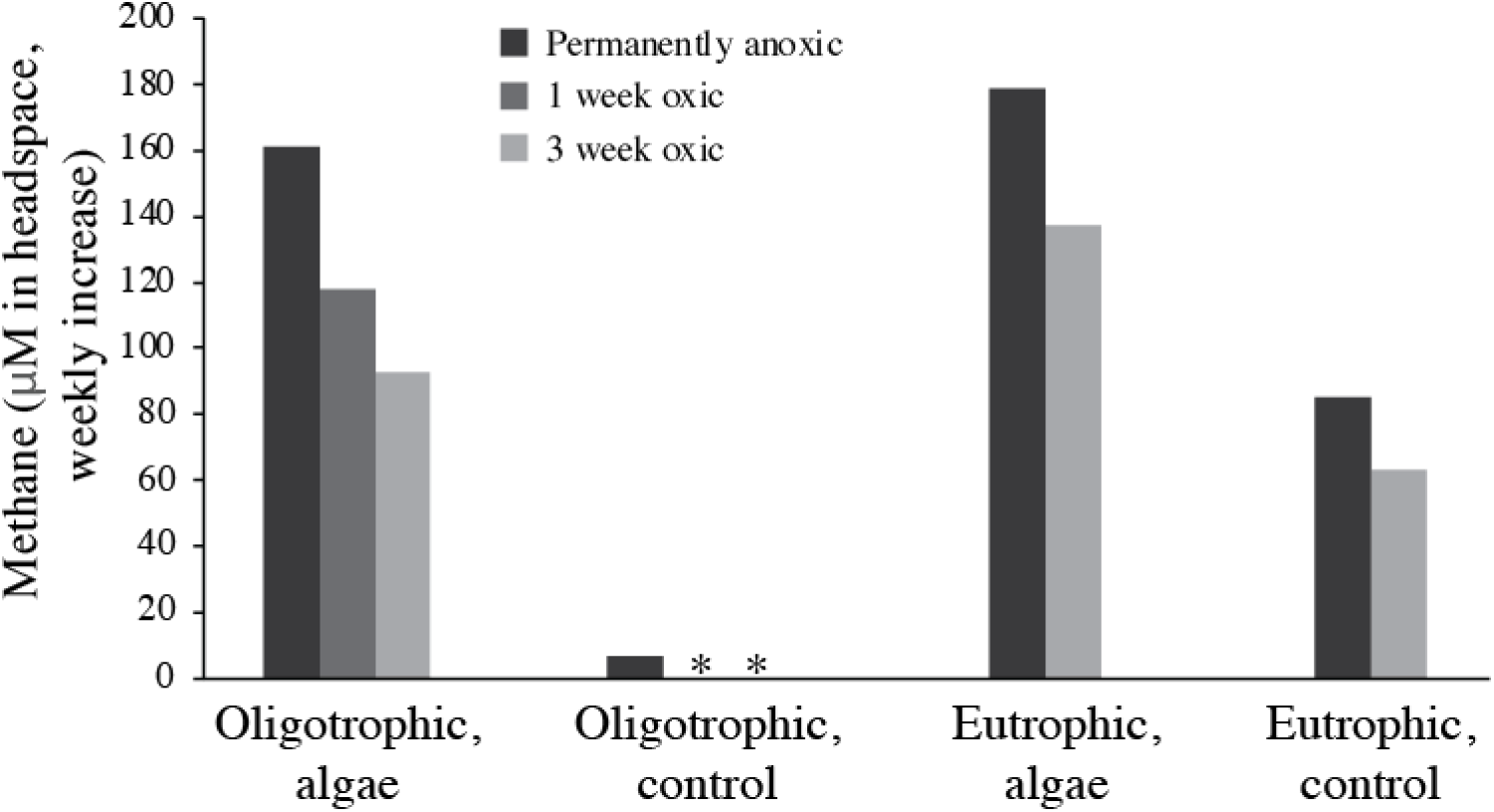
Weekly increase in headspace methane concentration in the oligotrophic and eutrophic slurry experiments, respectively, as derived from the linear phase of the methane concentration plots (Fig. 2). * no increase in concentration detected.

In both the eutrophic and oligotrophic sediments, methane emissions started directly in the first week after the start of the anoxic experiments (Fig. 2). After oxygen removal from the oxic slurries, thus re-establishing anoxic conditions, methane emission also started immediately. Net methane emission continued until week 13 in the anoxic oligotrophic slurries, and two weeks longer in both types of the oxic slurries, despite the 2 weeks difference between the start of methane emission in these two oxic incubations (Fig. 2). The methane concentration in the oligotrophic slurries plateaued and remained constant between weeks 13 or 15 to week 28, at a concentration of 1900 (anoxic), 1500 (1-week oxic) or 1200 (3-weeks oxic) μmol per L headspace. The methane concentration in the eutrophic slurries did not plateau, although two phases could be identified in the algae-fed slurries: A phase of rapid increase in [CH_4_] was observed from t_0_ until week 7 in the anoxic algae-fed slurries, and from week 3 to week 9 in the 3-week oxic algae-fed slurries (Fig. 2). After this initial phase, the methane emission rate stabilized at a similar rate as was observed in the control experiments.

**Fig. 2.**
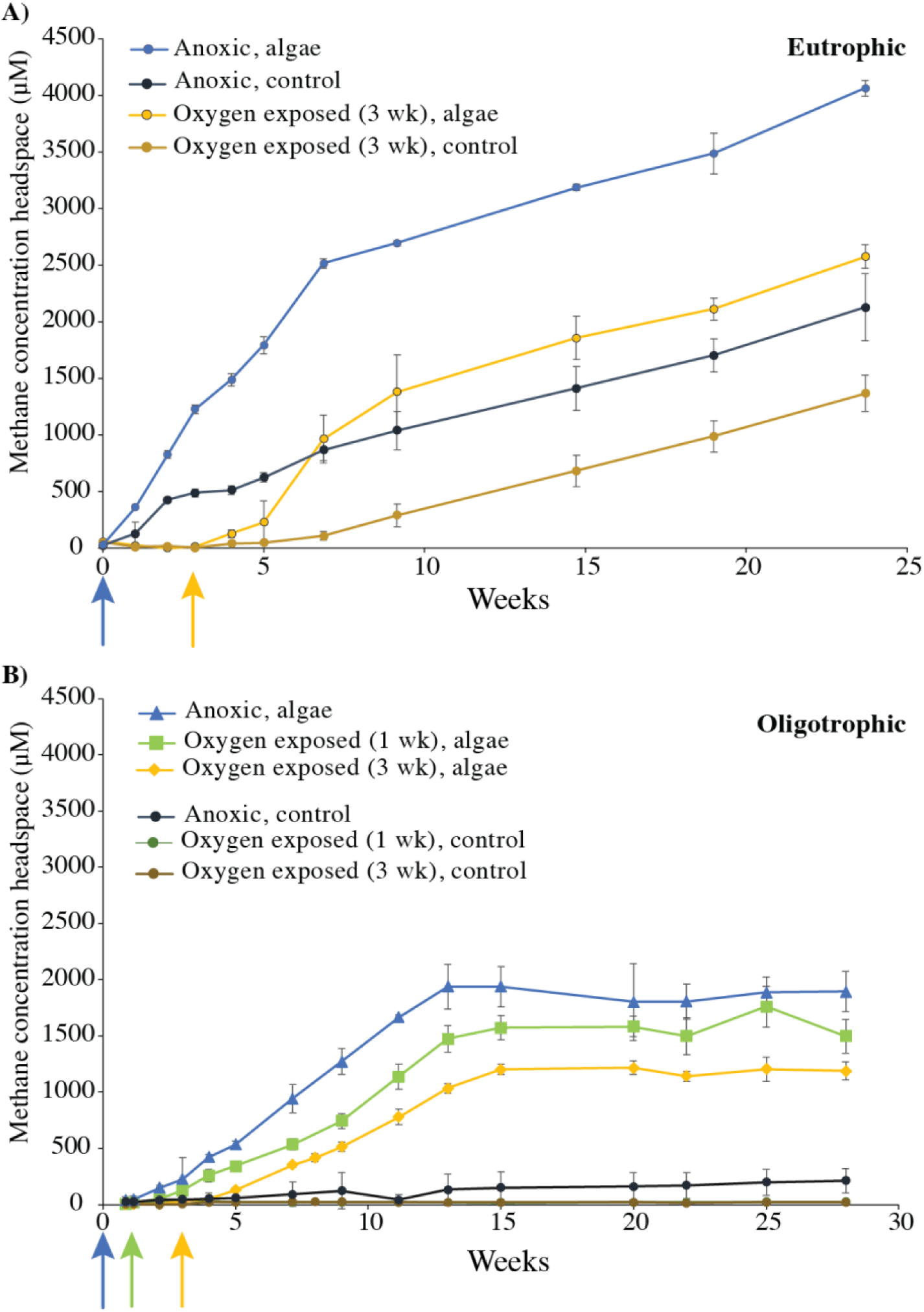
Methane concentration in the headspace of the slurry experiments with A) eutrophic and B) oligotrophic sediments. Arrows indicate the moment of oxygen removal, by flushing with N_2_, and the addition of anoxic sediments (see methods).

The weekly methane emission in the algae-fed oligotrophic sediments was significantly higher than in the control incubations, both under the permanently anoxic and temporary oxic conditions (Fig. 2). The background methane emission in the oligotrophic control experiments was below the detection limit in the initially oxic experiments, and around 9 μmol per week in the anoxic control sediments. The addition of algal biomass to the oligotrophic slurries increased the methane emission almost 25-fold under permanently anoxic conditions (to 161 μM per week), but only 17-fold and 14-fold under the 1-week and 3-week oxic conditions (118 μM and 93 μM per week), respectively (Fig. 1, Fig. 2).

The addition of algal biomass also increased the methane emission from the eutrophic slurries, although the presence of oxygen had as big an effect as the presence or absence of additional biomass. The background methane emission under anoxic control conditions was similar to that of the oxic conditions with additional algal biomass, resulting in a total amount of methane emitted of 2600 and 2100 μM for the oxic with algae and anoxic control, respectively, corresponding to an increase of only 20% in emitted methane (Table S1; Fig. 2). The methane emission rate was initially higher in the oxic incubations with algae, but because the phase of high emission was of a short duration (6 weeks duration, from week 3 to week 9), the total emission did not strongly exceed the total anoxic control emission. The high emission phase in the anoxic algae experiment was also short (t_0_ until week 7), but due to the high weekly emission rate of 180 μmol, the total amount of methane produced after 24 weeks was twice as high as the methane emission in the anoxic control experiment, and 1.5 times higher than in the oxic algae experiment (Fig. 2; Table S1).

Both the oligotrophic and eutrophic incubation experiments received equal amounts of algal biomass. The methane emission rate in the oligotrophic anoxic experiments increased from 6.9 to 161 μM per week due to the algae addition, whereas the eutrophic anoxic rate increased from 85 to 179 μM per week, showing a much larger increase in the oligotrophic experiments. The same holds for the 3-weeks oxic experiments, which increased in weekly rate from 0 (control) to 93 (with algae) in the oligotrophic experiments and from 63 to 137 in the eutrophic experiments, respectively. Due to the shorter duration of this high-rate methane emission the total methane produced as a result of the algal biomass was, however, similar (Fig. 2).

Experiments with whole sediment cores, rather than sediment slurries, showed a similar effect of oxygen exposure and algal biomass additions. Although the variation within the experiments with whole cores was much larger than in the more controlled slurry experiments, still a significant effect of oxygen exposure, and of algal biomass addition, was observed (Fig. 3).

**Fig. 3.**
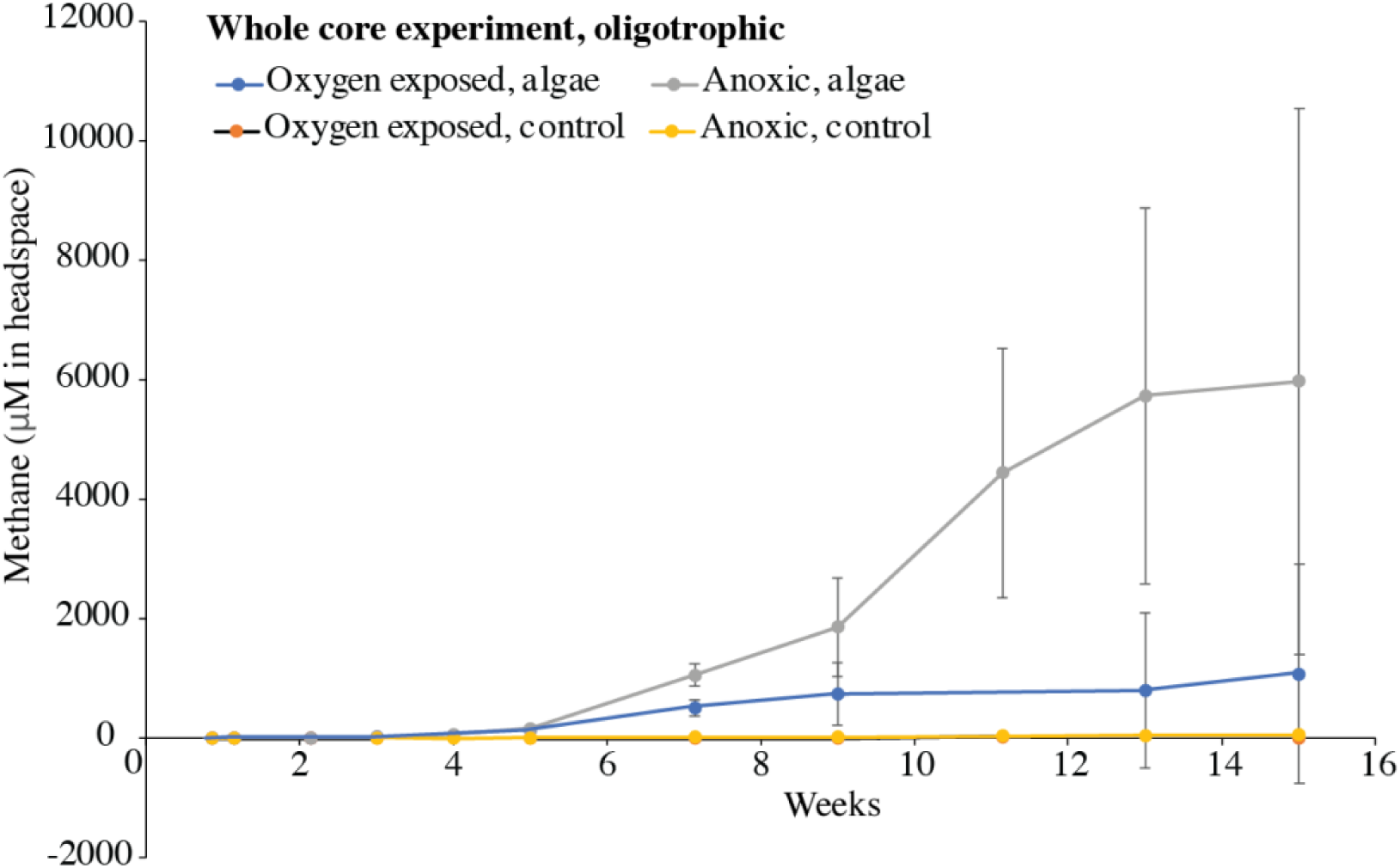
Methane concentration in the headspace of the whole core experiment under oxygen exposed (air headspace above overlying water) or anoxic (N_2_ headspace) conditions, with and without the addition of algal biomass, respectively. A cut-out of the lower values that highlights the methane concentrations at the start of the experiments is available in Fig. S6.

## Microbial community

### Methanogens

The relative abundance of methanogens in the eutrophic sediments was substantially higher than in the oligotrophic sediments. In both the oxic and anoxic eutrophic experiments, the relative abundance of OTUs assigned to methanogenic orders was mostly between 4 and 7% of the total 16S rRNA detected sequences, whereas in the oligotrophic incubations, this was between 0.5 and 3% for both the oxic and anoxic incubations (Fig. 4). Generally, both the algae and control experiments followed the same trend over time, as well as the oxic and anoxic experiments. In the eutrophic experiments, the oxic and anoxic experiments showed different trends in the first weeks, prior to the removal of oxygen. The anoxic experiments initially peaked in methanogen abundance (6 – 7%), whereas the abundance in the oxic experiments was initially low (1.5 – 2.5%). After 4 weeks, all experiments established around relatively stable values in the range of 4 – 7% (Fig. 4). Thus, a slight increase in the relative abundance of methanogens over time was observed in both the oligotrophic and eutrophic incubations.

**Fig. 4.**
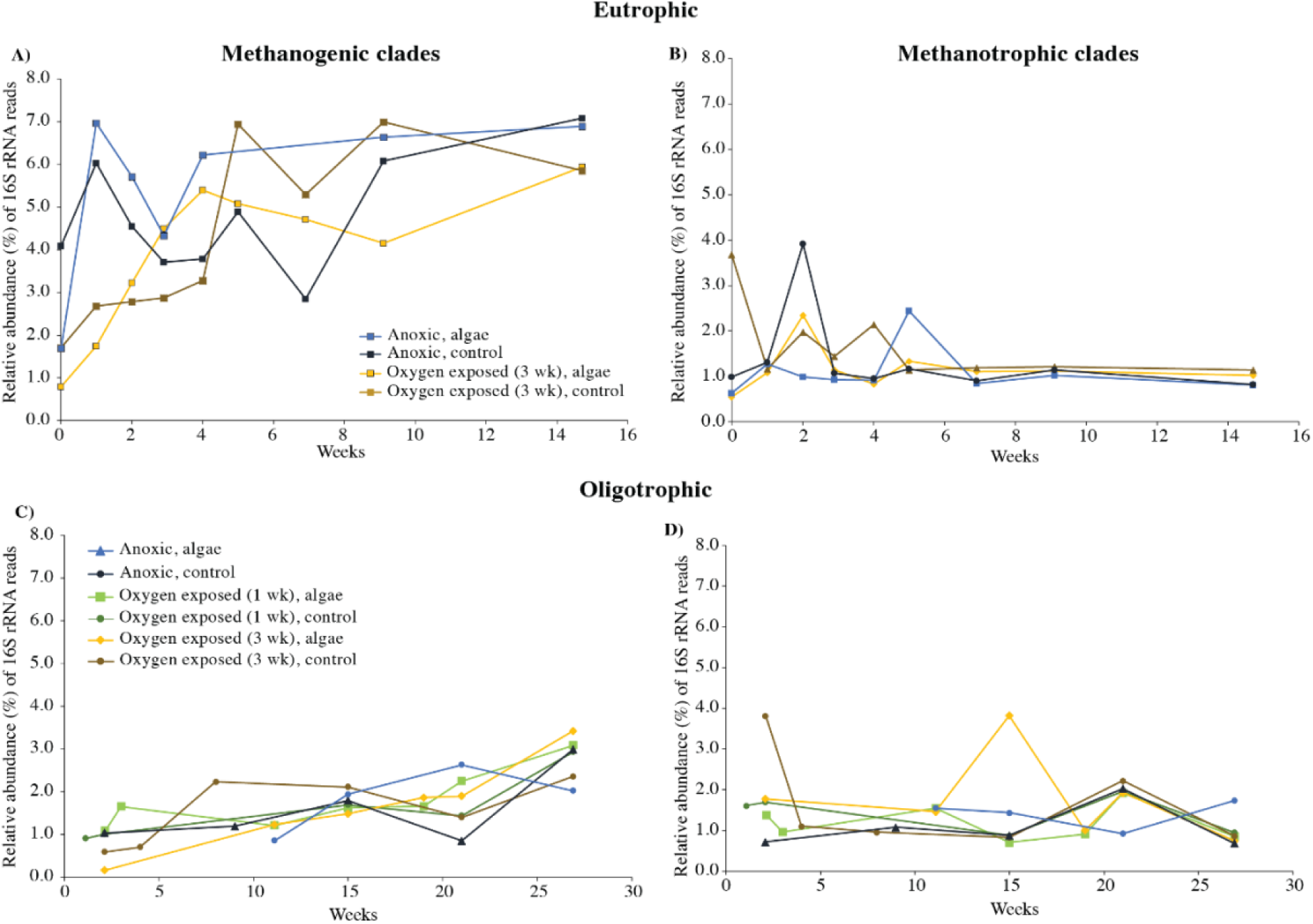
Relative abundance of OTUs assigned to known methanogenic (A; eutrophic, C; oligotrophic) and methanotrophic (B; eutrophic, D; oligotrophic) clades, in the slurry experiments. Note that the oxic experiments received additional sediments, of the 5 – 15 cm depth interval, after the initiation of the anoxic conditions, after 1 or 3 weeks.

The methanogenic community was dominated by OTUs assigned to the order Methanomicrobiales (Fig. S4). The contribution of Methanomicrobiales to the total 16S rRNA sequences assigned to methanogenic orders was, however, higher in the eutrophic (average of 75%) than in the oligotrophic experiments (average of 65%). Methanomassiliicoccales made up a larger fraction in the oligotrophic experiments (average of 29%, versus 12% in eutrophic experiments). Methanosarcinales were relatively more abundant in the eutrophic experiments (Fig. S4). In the oligotrophic experiments, no patterns were observed over time, nor differences between treatments. In the eutrophic experiments, Methanomicrobiales dominated both the oxic and anoxic experiments. However, in the permanently anoxic incubations, more OTUs are assigned to Methanosarcinales, whereas in the temporarily oxic incubations, more OTUs are assigned to the order Methanomassiliicoccales.

### Methanotrophs

In both the oligotrophic and eutrophic incubations, the relative abundance of methanotrophs belonging to the order Methylococcales was around 1 – 2% for all treatments. Methanotrophs belonging to the order Methylomirabilis were detected in all treatments, with a relative abundance around 0.5% in the oligotrophic incubations and 0.2% in the eutrophic ones (Fig. 4). No difference in the relative abundance of methanotrophs was observed between treatments, and no trend was observed over time.

The majority of OTUs within the Methylococcales order were assigned to the genus *Crenothrix* in all eutrophic treatments (20 – 99% of Methylococcales reads), followed by the genus *Methylobacter* (0.9 – 30.6% of *Methylococcales* reads, Fig. S5). However, recent research on the SILVA annotation within the Methylococcales order has shown that a differentiation between *Crenothrix* and certain groups of *Methylococcaceae* cannot be supported, and the weight of the differentiation between these two groups is thus limited, currently (van Grinsven et al. 2022). At specific time points, the genus *Methylomonas* showed particularly high peaks in its relative abundance (up to 74% of Methylococcales reads) in both the eutrophic and oligotrophic incubations, driving down the relative contribution of other groups, but this was not consistent over time (Fig. S5). The methanotrophic community in the oligotrophic experiments was less dominated by sequences assigned to “*Crenothrix*”, and rather than a higher abundance of *Methylobacter* assigned OTUs (Fig. S5) and a higher abundance of genera that were marginal in the eutrophic incubations, such as *Methyloparacoccus*. However, overall the methanotrophic communities looked relatively similar and no trends could be established over time, nor differences between the oxygen or algae treatments (Fig. S5).

### Volatile fatty acids

The concentration of different volatile fatty acids (VFAs) was traced in the oligotrophic slurry experiments, during the first 8 weeks, as shown in Fig. 5. Acetate was the dominant VFA, with concentrations 10-100x higher than the other VFAs. In general, VFA concentrations were highest in the anoxic algae incubations, although the concentrations of formate and pyruvate were equally high, or higher, in the 1-week oxygen exposure experiments. VFA concentrations in the 3-week oxic experiments were much lower, at all timepoints and for all VFAs.

**Fig. 5.**
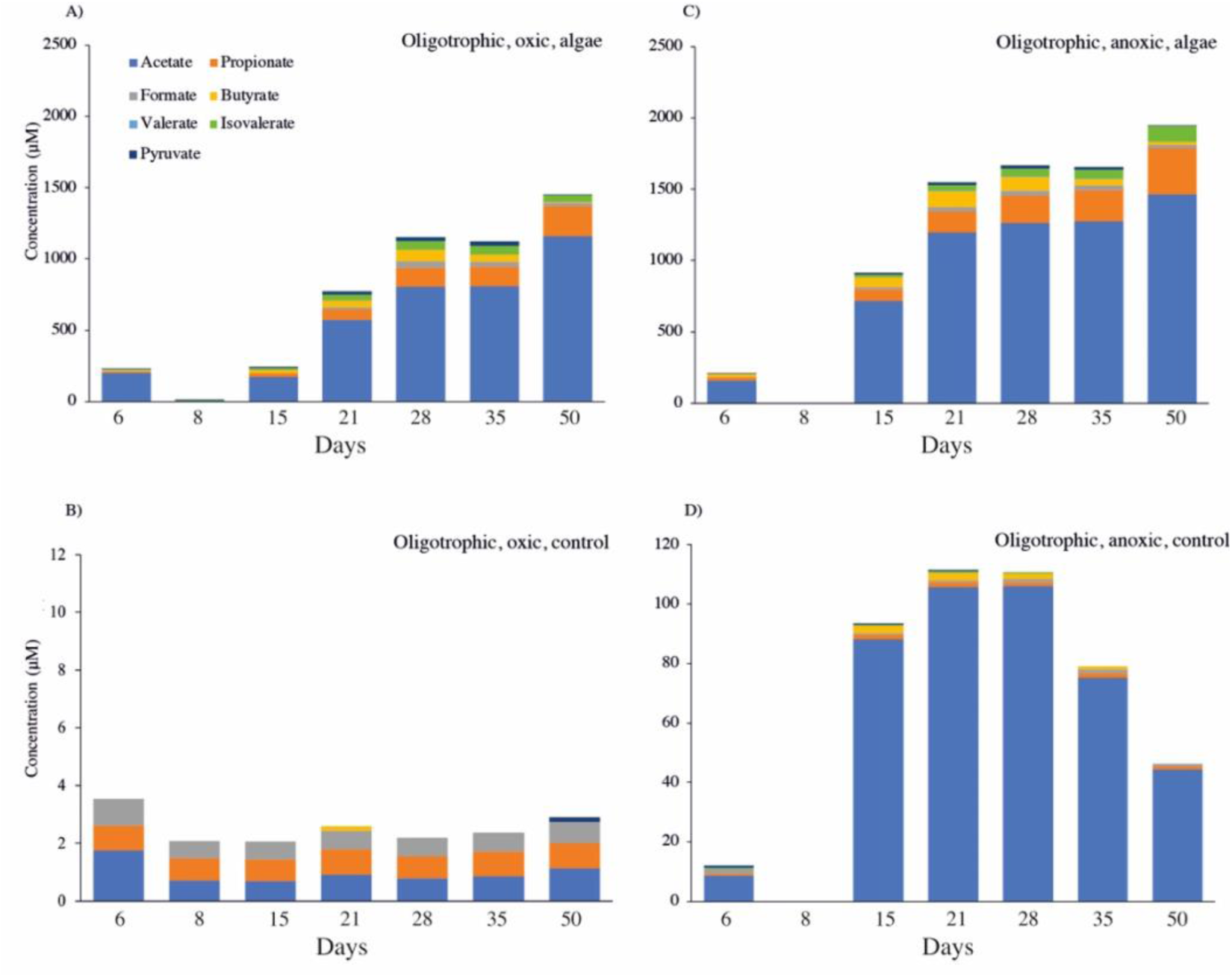
Volatile fatty acid concentrations during the first 8 weeks of the 1 week oxic (A, B) and anoxic (C, D) oligotrophic experiments. Note the different y-axes. Each bar represents the average of triplicate samples at each timepoint. The samples of day 8 of the oxic experiments were taken shortly after anoxic conditions were created. No samples were taken on day 8 of the anoxic experiments.

Acetate and propionate concentrations were still increasing after 8 weeks, whereas butyrate, isovalerate, formate, and pyruvate concentrations were decreasing after peak values during weeks 3 through 5. The VFA concentrations in the control experiments, without algal biomass additions, were <2 μM for each component, except for the acetate concentrations in the anoxic control experiment (Fig. 5A).

## Discussion

### Trophic state effects

Eutrophication of lakes is an ongoing, and increasing, problem. Our results show a major legacy effect based on the eutrophication state of the lake, similarly to earlier studies on these lakes (Fiskal et al. 2019; Han et al. 2020).Our control experiments show that even after nutrient and carbon inputs are completely stopped, the continuous background methane emission is much higher in eutrophic sediments than in oligotrophic sediments: The methane emission was 12 times higher in control experiments with eutrophic sediments than oligotrophic sediments, also though no fresh material input was delivered over 160 days (Fig. 1, Table S1). The TOC concentrations in sediments of Lake Lucerne (oligotrophic) and Lake Baldegg (eutrophic) are comparable, according to (Fiskal et al. 2019), but the sedimentation and TOC deposition rate is much higher in the eutrophic Lake (Fiskal et al. 2021).

These legacy effects are, however, contrary to our expectation of little influence on the conversion of fresh biomass to methane. The response to the input of new algal material was similar in magnitude in both oligotrophic and eutrophic sediments (Fig. 1; Fig. 2). Possibly, the large amount of non-degraded organic carbon that was added, led to a shortage in electron acceptors other than CO_2_. In both types of sediment, methane emission started directly after the addition of the substrate (Fig. 2). Both this study and earlier studies on these lakes, showed an initial population of methanogens was present in the sediments (Fig. 4; Meier, van Grinsven et al., *in prep*). Although weekly emission rates were higher in the eutrophic sediments, the total amount of methane emitted as a result of the algae input was comparable (± 2000 μM; Table S1). We pinpointed the end of the biomass addition effect at the timepoint when the oligotrophic sediments reached a stable [CH_4_] level (week 13 – 15) and when the eutrophic sediments reached a weekly emission rate that was similar to that of the control experiment (week 7 – 9; Fig. 2).

The continuous high methane emission in the eutrophic sediments (weekly increase of 85 μM in control experiments), in contrast to the low methane emission in the oligotrophic sediments (weekly increase of 7 μM in control experiments), leads to the expectation that a more abundant and more active methanogenic microbial community may exist in the eutrophic sediments. Although the methanogenic community was indeed higher in its relative abundance (3 – 7%) in the eutrophic experiments than in the oligotrophic sediments (0.5 – 4%), the methanogenic community was comparable in its CH_4_ emission response to fresh biomass in both types of sediments. The fact that no lag phase was observed in the methane emission response in the oligotrophic sediments, supports the idea that there was already an abundant methanogenic community present that directly reacted to the additional substrate. It is surprising that the oligotrophic methanogenic community is able to respond this rapidly to a high amount of incoming algal material, as such high pulses of organic material are uncommon in oligotrophic Lake Lucerne (Fiskal et al. 2019). The main methanogens present, belonging to the orders Methanomicrobiales and Methanomassiliicoccales, are known for doubling times in the order of hours to days, under ideal conditions. They might therefore have responded to the additional substrate input by either increased per cell methane production rates, or by rapid community growth. No strong increase in relative abundance of methanogens was observed in the oligotrophic algae-fed experiments (Fig. 4c). An increase in methanogenic cell numbers may have been masked by an increase in the total microbial community, as relative abundances are reported here. An alternative explanation would be that not the cell numbers, but the per cell methanogenesis rate, increased in the methanogenic community of the oligotrophic sediments, following the algal biomass inputs.

### Oxygen pulses

The effect of oxygen penetration depth on methane emission from lake sediments is well established. However, these studies generally address stable oxygen conditions, in the seasonal to kyear range ((Huttunen et al. 2006; Sobek et al. 2009). Here, we look at short oxygen pulses, as a potential mediative measure for lakes with anoxic bottom water. The presence of oxygen for a short, 3-week period at the start of the incubation had major implications for methane emissions over the course of the entire experiments. The total release of methane was significantly lower in the treatments that had experienced an oxic period (Fig. 1; Fig. 2, Table S1). Most likely, part of the algal biomass was converted to CO_2_ during the oxic period, and was therefore not available for methanogenesis anymore. This is supported by the peak in CO_2_ emissions that was observed during the oxic period of the experiments (Fig. S7; Fig. S8). However, due to difficulties in translating headspace CO_2_ concentrations to dissolved CO_2_, it is not possible to make a carbon mass balance, to see how much is indeed released as CO_2_. Part of the produced CO_2_ will also be converted prior to release to the headspace, leading to underestimates that cannot be sufficiently quantified. As CO_2_ has a much lower warming potential per mole than methane (approximately 28 times lower on a hundred year basis, (Forster et al. 2021) the release of CO_2_ is strongly preferred over that of methane in light of global warming. Besides CO_2_, part of the carbon may have been converted to microbial biomass during the oxic period, and is stored as such in the sediments. (Sobek et al. 2009) published a weak linear relationship between the diffusive methane flux from lake sediments, and the oxygen penetration depth at those locations. A direct comparison with this study is, however, difficult to make, as there are likely other factors involved that affect both the oxygen penetration depth and the methane production, such as carbon content of the sediments.

No methane release was measured during the oxic period in these slurry experiments, from none of the treatments. It is likely that little to no methane was produced in the sediments. Most methanogens are known to be strict anaerobes, that are not active under oxic conditions. However, the tolerance of methanogens to oxygen is debated, and additionally differs between clades (Kiener and Leisinger 1983; Zinder 1993; Kato et al. 1993). The experiments were setup with only the top 5 cm of the sediment cores, both in the eutrophic and oligotrophic incubations, to not expose the deeper sediments to oxygen, and to stay closer to natural conditions mimicking artificial aeration. During the oxic incubations, anaerobic micro-niches were most likely present, as these are common in water-logged, particle-heavy media such as sediments (Jørgensen 1977), and these niches may have harbored active methane producing communities. However, methane production in such niches was not high enough to be detected as a methane concentration increase in the headspace. The produced methane would likely also be rapidly consumed by methanotrophs. These were present according to the 16S rRNA sequencing and can thrive under interchanging oxic and hypoxic conditions (van Grinsven et al. 2020). Directly after oxygen was removed from the incubation bottles and sediments from 5 – 15 cm depth were added, methane started to build up again (Fig. 2). Aerobic methanotrophy was disabled due to the removal of oxygen, and anaerobic methanotrophs may have been electron acceptor limited under the created anoxic conditions. The immediate buildup of methane in the headspace after the creation of anoxic conditions, suggests that algal biomass was directly converted to methane precursors, or those precursors were present in the incubation bottles from the oxic period. The concentration of VFA was lowest at the moment directly following the oxic-anoxic switch and then increased (Fig. 5). A dilution effect by the addition of extra sediments (see *Methods*) artificially lowered the concentration compared to the timepoint directly before the oxic-anoxic switch. However, from the comparison between the control experiments under both oxygen-pulse and anoxic conditions, it is clear that the effect of the oxic period is large: VFA concentrations are 10x higher in the permanently anoxic control experiment (Fig. 5B and D).

A similar effect of an oxic-anoxic switch was observed by (Frenzel et al. 1990), who observed an abrupt increase in sedimentary methane emissions when the oxygen concentration in the water overlying their core experiments dropped below 18 μM. They assigned the difference between oxic and anoxic methane emissions solely to an increased activity of methanotrophs under oxic bottom water conditions.

Although there was still an active methanogenic community after the oxygen exposure, a lasting effect was observed: the weekly methane emission rate in the oxic incubations in the oligotrophic experiments, after oxygen removal, did not reach the same values as the permanently anoxic conditions. There was also a significant difference in weekly emission rate between the 1-week and 3-week oligotrophic oxygen exposure treatments (Fig. 1, Table S1). This effect was not only visible in the methane emission rates and total methane emission, but also in the VFA concentrations (Fig. 5). This could partly be attributed to the fact that a fraction of the algal biomass had been converted to CO_2_ instead of methane prior to the oxic-anoxic switch, but it seems the processes converting OM to methane, including intermediated steps, has been affected: The rate with which algal biomass is converted becomes slower and less efficient after an oxygen exposure period, as can be seen by the lower per-week rates in the post-oxic phase in Figure 2 and Table S1. However, a lower per-week increase in [CH_4_] in the headspace may be either the result of lower methane production or a higher consumption by methanotrophy.

Stable isotope analysis of the headspace methane in the stable, post-algal biomass degradation phase (17 weeks of oligotrophic, and 11 weeks of eutrophic experiments, Fig. 6), showed more negative δ^13^CH_4_ values in the algal biomass experiments. The δ^13^C signal of the algal biomass likely decreased the δ^13^CH_4_ values in the algal addition experiments, with a larger effect in the oligotrophic lake, where the relative contribution of algal biomass was largest, compared to the organic matter already present in the sediments. When comparing the oxic and anoxic experiments, only the oligotrophic experiment showed significant differences: the δ^13^CH_4_ values were lower (more depleted) in the anoxic than in the oxic incubations, both with and without algal biomass additions. This could be caused by differences in methanogenesis pathways, e.g. methane production from CO2 yields more 13C-depleted methane than acetoclastic methanogenesis (Conrad 2005)). As no changes in the methanogenic community were observed between the oxic and anoxic oligotrophic treatments, it is unlikely that a change in the community caused the dominant methanogenesis pathway to swap and to cause the differences in the δ^13^CH_4_ values. A further explanation is that differences in rates of methanotrophy caused the observed changes in ^13^C-compositions of methane. Indeed, increased rates of methanotrophy under oxic conditions would be expected to contribute to a less depleted isotopic composition of the remaining methane (Barker and Fritz 1981).

**Fig. 6.**
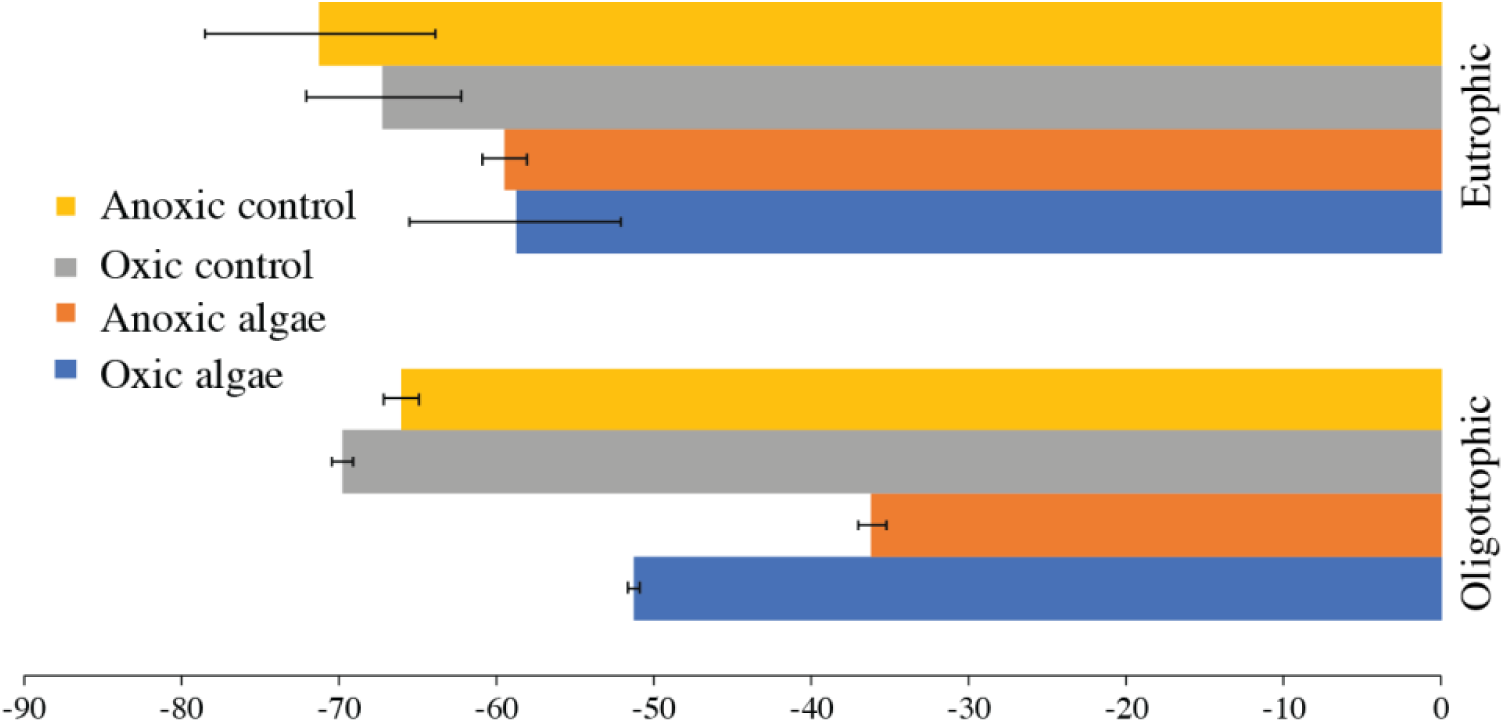
δ^13^CH_4_ values of headspace methane, after 11 weeks (eutrophic sediments) or 17 weeks (oligotrophic) of slurry incubations.

The VFA data shows a generally lower concentration of VFAs in the oxygen-exposed incubations, but not a different composition of the total VFAs (Fig. 5).

### Methane emissions and implications

The methane release from the sediments was linear in all slurry treatments. No initial lag, nor logarithmic start was observed in any of the treatments, nor a gradual decrease in methane release rate over time (Fig. 2). In all algal-fed slurry experiments, two linear phases with different coefficients could be observed (Fig. 2). Although the algal biomass is expected to consist of a combination of more and less labile compounds, no higher initial degradation was observed, suggesting that the conversion of algal biomass to methane was not dependent on specifically highly labile compounds. The whole core experiments surprisingly also did not show a lag phase. Both in the oxic and anoxic algal-addition experiments, methane emission started immediately (Fig. 3; Fig. S6). This is surprising given the fact that methanogenesis is not expected to occur in the surface sediments, but rather deeper into the sediments (from 3 cm depth onwards, as modeled for Lake Lucerne by (Fiskal et al. 2019)). The transport of carbon towards this deeper methanogenic layer, could be expected to take time, and therefore, a lag phase between algal addition and methanogenesis from these compounds, could be expected. This was, however, not the case: degradation at the sediment surface apparently directly results in methane emissions (Fig. 3; Fig. S6).

Generally, only the sediment surface is affected by the oxygen conditions in the bottom water; deeper sediments are anoxic, due to the low diffusion coefficient through sediments. Maerki *et al*. (2009) investigated the oxygen, carbon and nitrogen dynamics of lake sediments, and stated that short term (weeks to months) oxygen exposure is insufficient to change the reactivity spectrum of eutrophic Lake Zug sediments, that the exposure times are too short for that. Our whole core experiment shows, however, that despite the fact that the methanogenic layer is deeper in the sediments than the bottom water affected layer, the conditions in the bottom water are still of key importance for the methane emissions from the sediments following the deposition of algal material, for example after an algae bloom in the surface waters. (Maerki et al. 2009) also state that over 95% of the anaerobic mineralization in Lake Zug sediments was due to methanogenesis, and that methane oxidation was responsible for over half of the oxygen consumption at the sediment surface. If a similar situation is the case in our eutrophic Lake Baldegg, changes in the methane cycling are likely to have substantial effects on the carbon and oxygen cycling in the shallow sediments.

Our experiments show that the effects of a short (1-3 week) oxygen exposure can last for several months, i.e. decreasing methane emissions without changing the methane-related microbial community (Fig. S4, Fig. S5). We believe these findings should be further explored in environmental settings. If brief pulses of oxygen, like the 1- and 3-week oxygen exposure periods tested here, have the capacity to reduce longer-term methane emissions, we believe this could be promising, especially if applied directly after an algal bloom, as tested here. A slower release of methane from the sediments to the water column could also increase the potential of water column methane oxidation, as more time is available for the process of oxidation and thus removal of methane. Such an effect could further limit the emissions of methane at the water-air interface, and thus to the atmosphere. The transfer of our results to field applications needs more research.

Artificial oxygenation is already applied in specific Swiss lakes. Previous research on the effect of artificial aeration on sediment processes indicated little effect on the sedimentary biogeochemistry, mainly due to the small effect on bottom water oxygen concentrations (Horppila et al. 2015); thus indicating that care should be given to effective bottom water aeration, as our study shows that enhanced oxygen concentrations in the bottom water will affect the methane dynamics of the sediments. Given the expectations of ongoing eutrophication in the upcoming decades, plus the global warming of lakes that further draws down oxygen levels, we believe this should be a topic for further research.

## Author contribution statement

Conceptualization by SvG, MAL and CJS. Original draft preparation by SvG, review and editing by SvG, NM, CG, MAL and CJS. Investigation and Methodology by SvG, NM and CG.

## Data availability statement

Raw reads of the 16S rRNA sequencing data is deposited and made publicly available in the online repository NCBI SRA, under accession number XXX (in progress).

## Acknowledgements

The authors thank Patrick Kathriner, Karen Beck, Kathrin Baumann, Cameron Callbeck and Dimitri Meier for help in the field and the lab.

The authors have no conflict of interest to declare.

## Supplemental information

**Table S1.** Methane emission from the slurry experiments.

Total methane emission
| Oligotrophic sediments |  |  | Eutrophic sediments |  |  |
| --- | --- | --- | --- | --- | --- |
|  | [CH<sub>4</sub>]<sup>a</sup> | Standard deviation |  | [CH<sub>4</sub>]<sup>b</sup> | Standard deviation |
| Permanently anoxic, algae | 1893 | 178 | Permanently anoxic, algae | 4063 | 68 |
| 1 week oxic, algae | 1497 | 147 | 3 week oxic, algae | 2576 | 105 |
| 3 week oxic, algae | 1186 | 81 |  |  |  |
| Permanently anoxic, control | 211 | 108 | Permanently anoxic, control | 2127 | 293 |
| 1 week oxic, control | 23 | 25 | 3 week oxic, control | 1367 | 162 |
| 3 week oxic, control | 19 | 7 |  |  |  |

a. Concentration in headspace in $\mu\text{M}$ after 28 weeks (constant since 14 weeks)
b. Concentration in headspace in $\mu\text{M}$ after 24 weeks

**Fig. S1.**
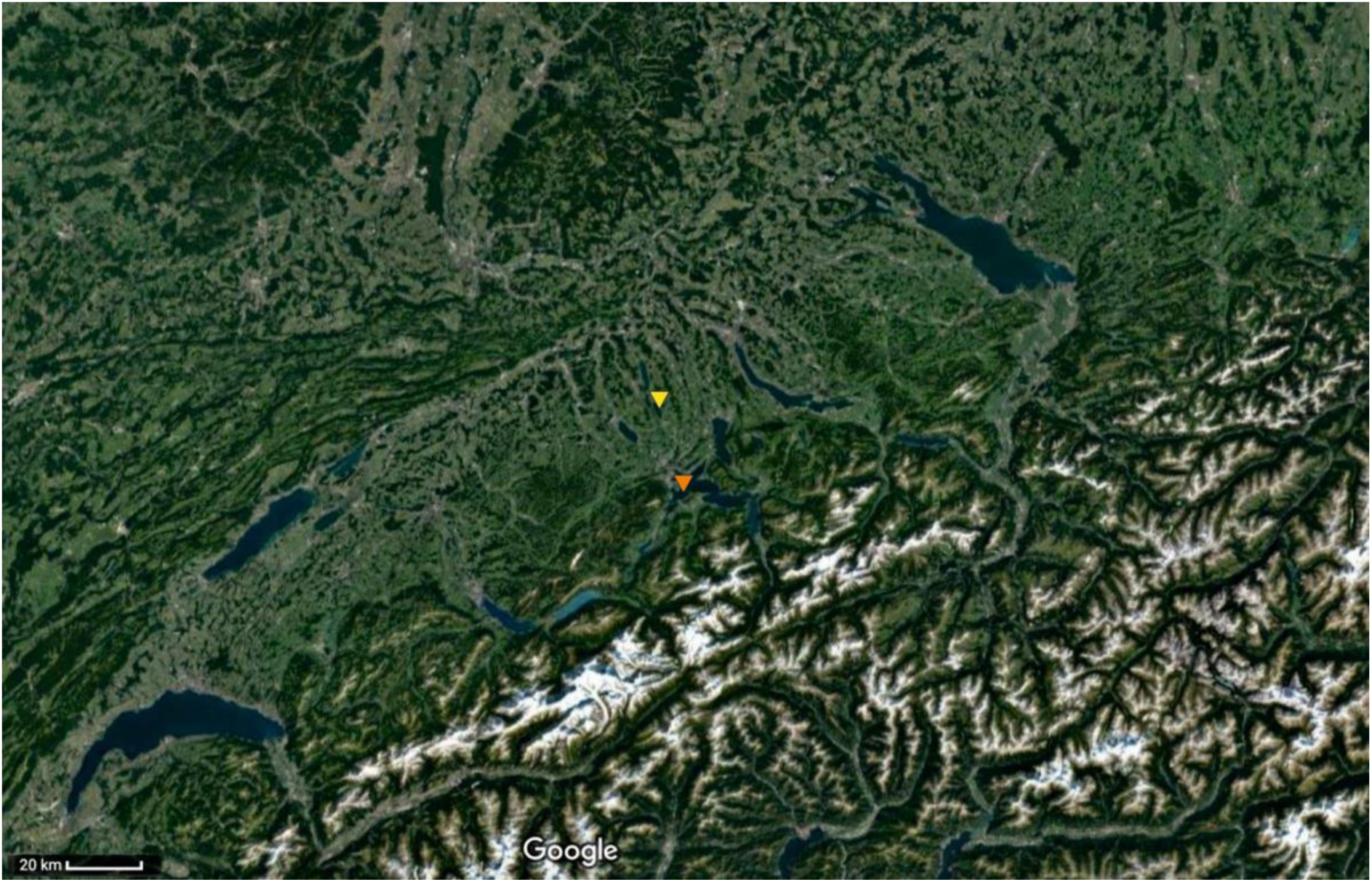
Map of the location of oligotrophic Lake Lucerne (orange) and eutrophic Lake Baldegg (yellow) in Switzerland, central Europe.

**Fig. S2.**
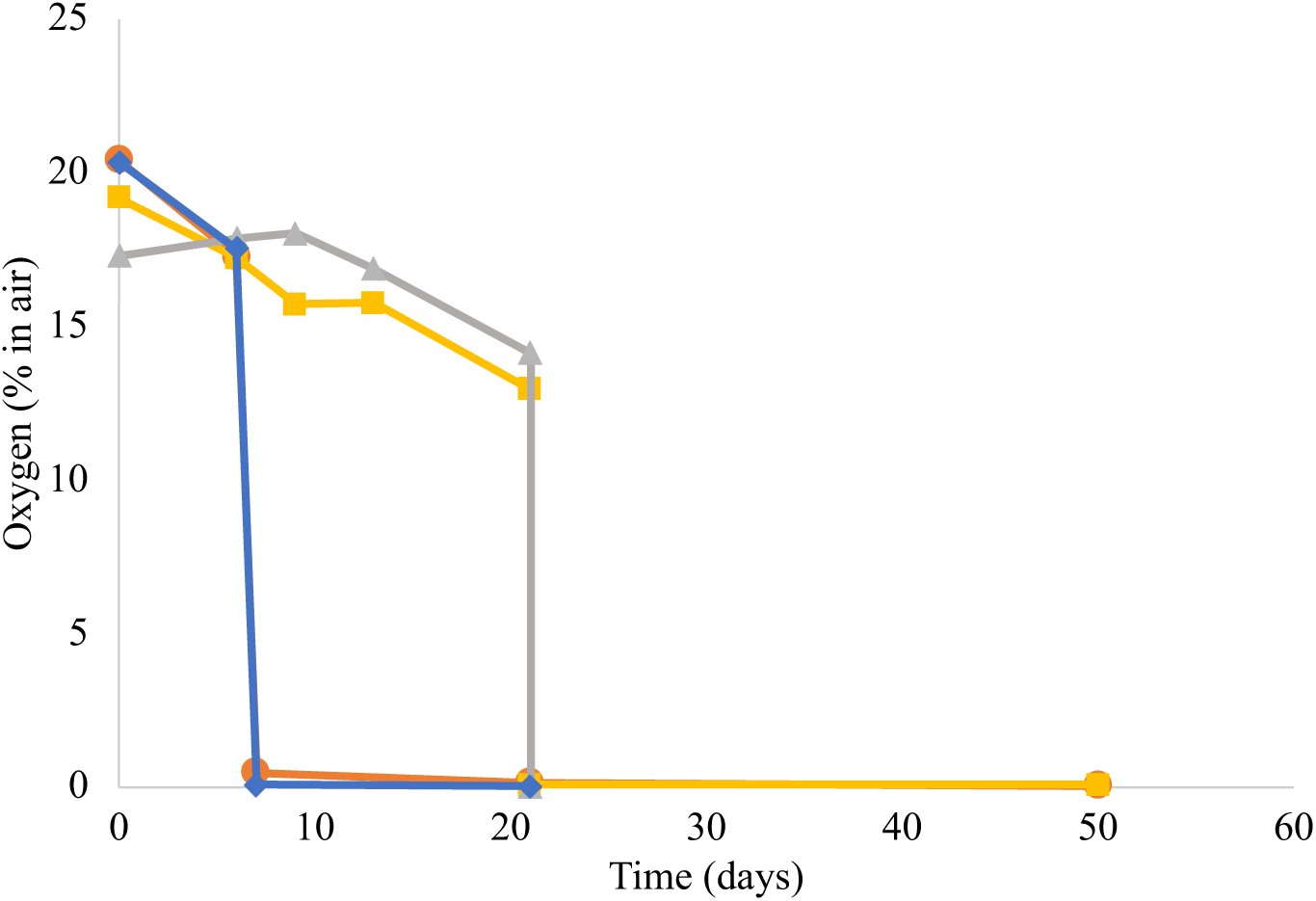
Oxygen concentration (in % of ambient air) in the slurry experiments. Blue and orange: two replicate bottles of the 1-week oxygen exposure experiment. Yellow and grey: two replicates of the 3-week oxygen exposure experiment.

**Fig. S3.**
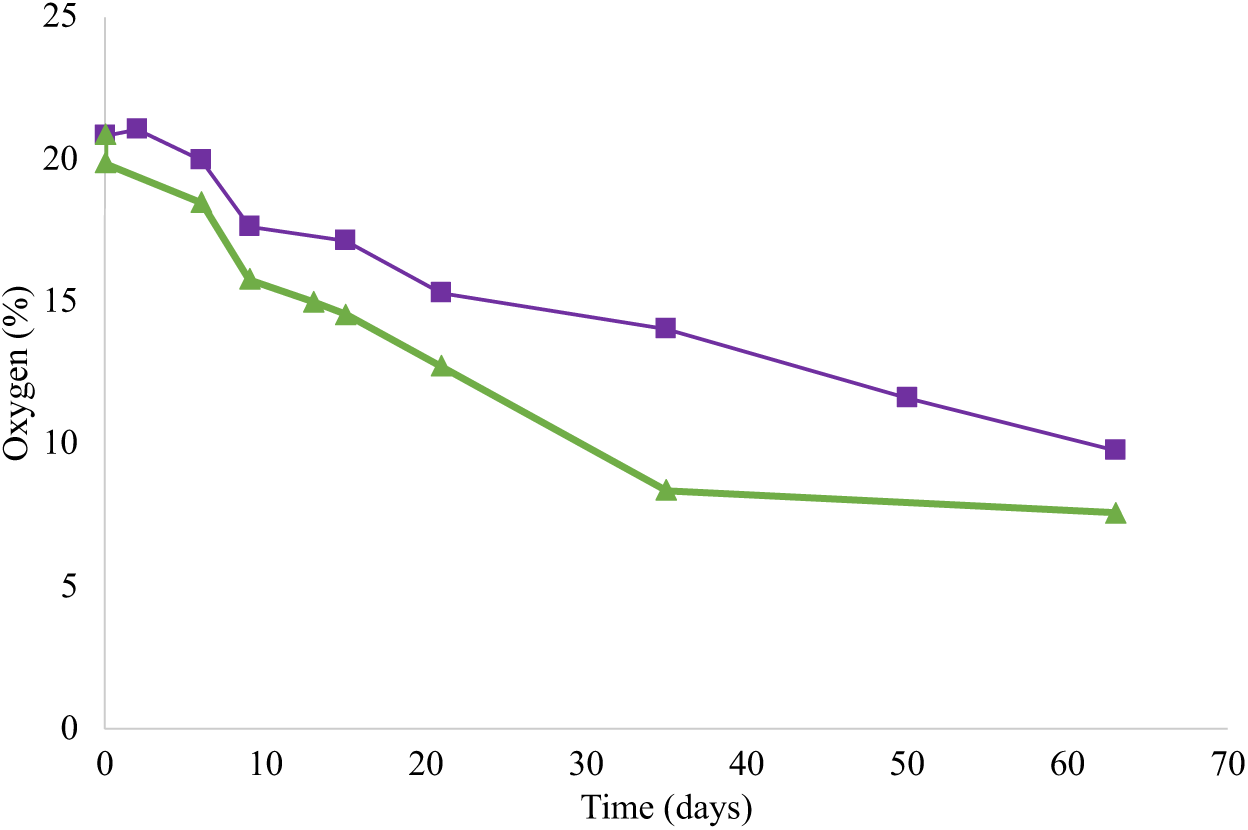
Oxygen concentration (in % of gas headspace) in the whole core experiments. Green: oxic algal biomass addition experiment, purple: oxic control experiment.

**Fig. S4.**
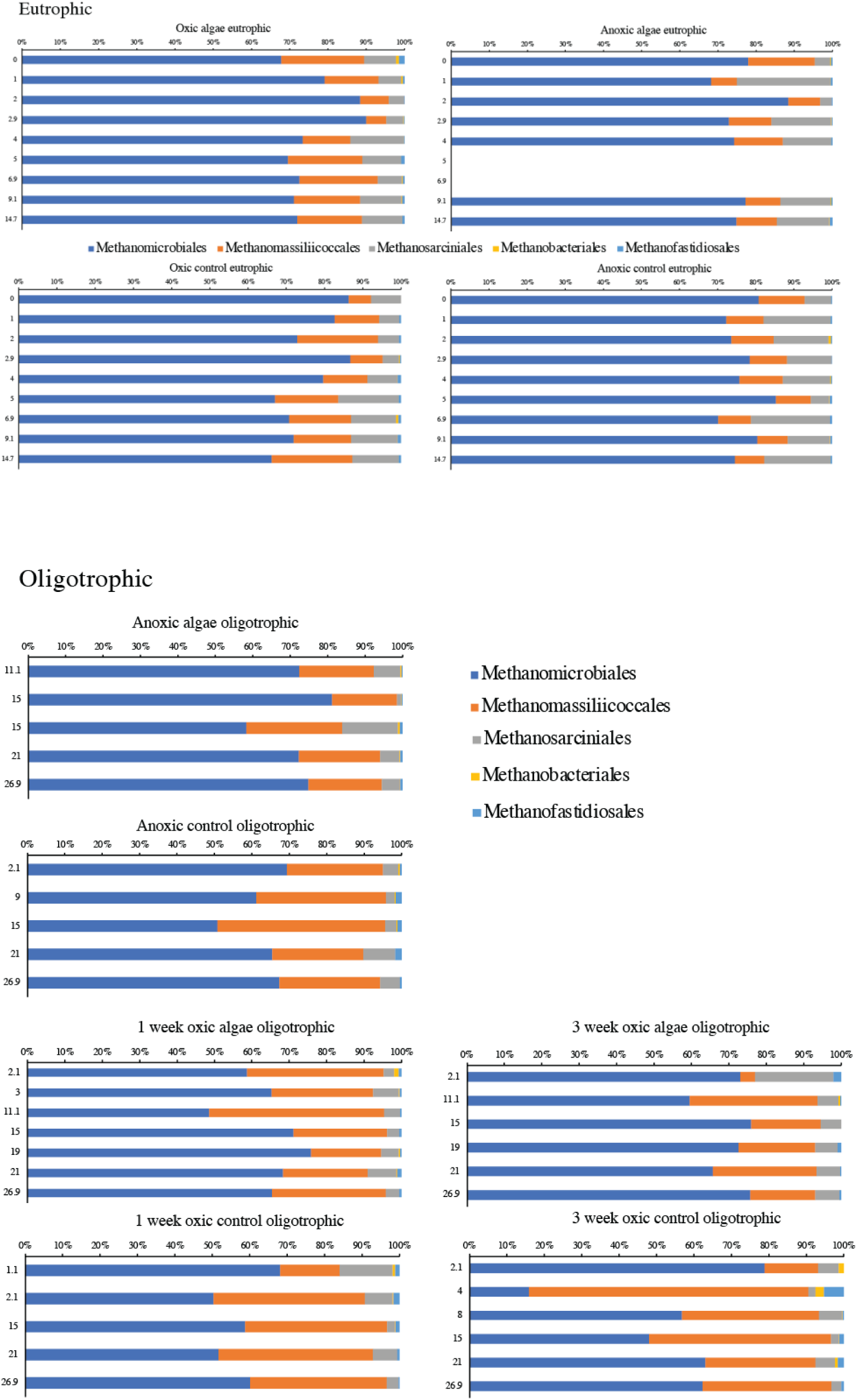
Distribution of OTUs assigned to methanogenic orders over the five most abundant orders for both the eutrophic and oligotrophic experiments. The y-axis shows the time in weeks since the start of the experiment.

**Fig. S5.**
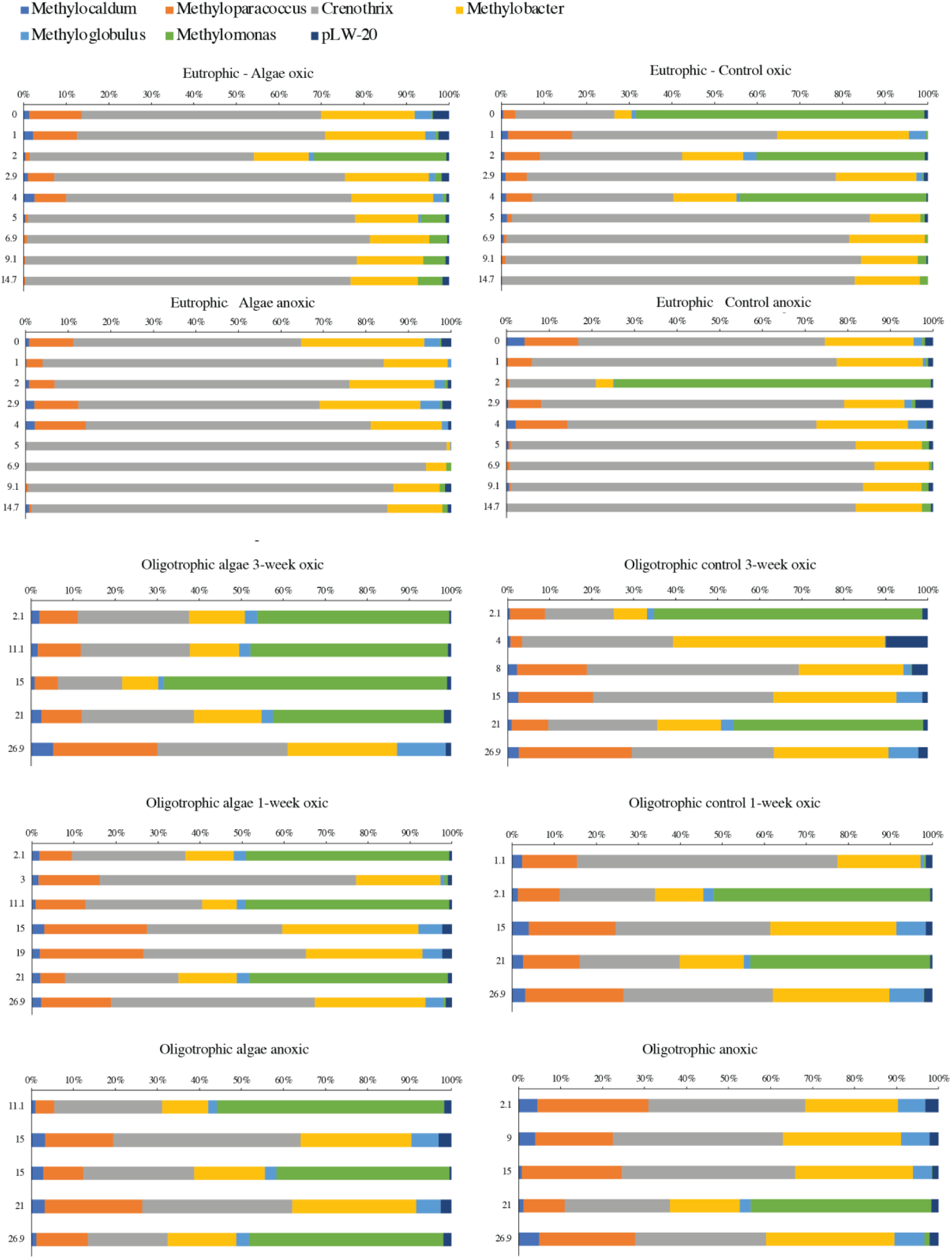
Distribution of OTUs assigned to methanotrophic orders over the most abundant orders for both the eutrophic and oligotrophic experiments. The y-axis shows the time in weeks since the start of the experiment.

**Fig. S6.**
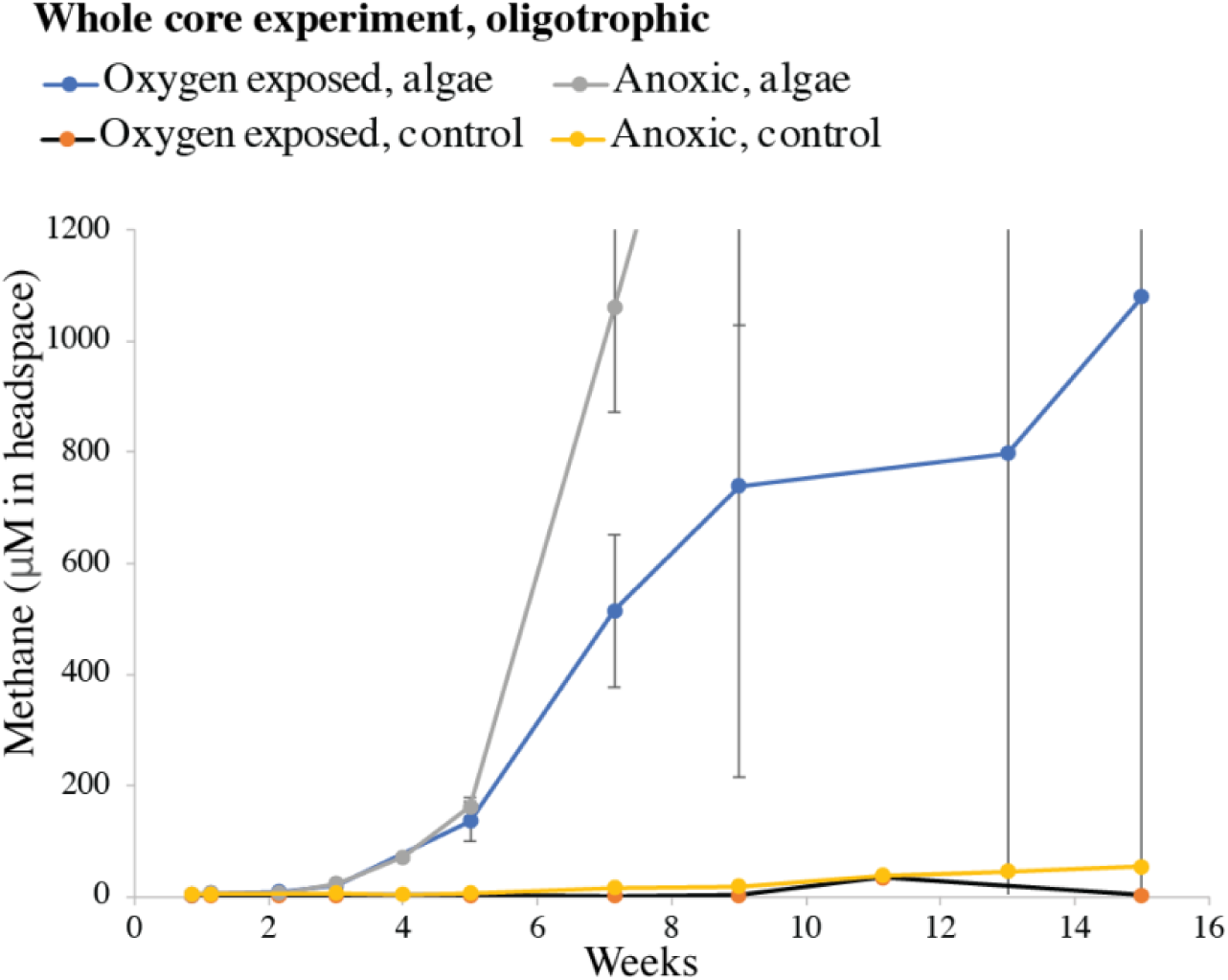
Cut-out image of Fig. 3, highlighting the lower part of the y-axis.

**Fig. S7.**
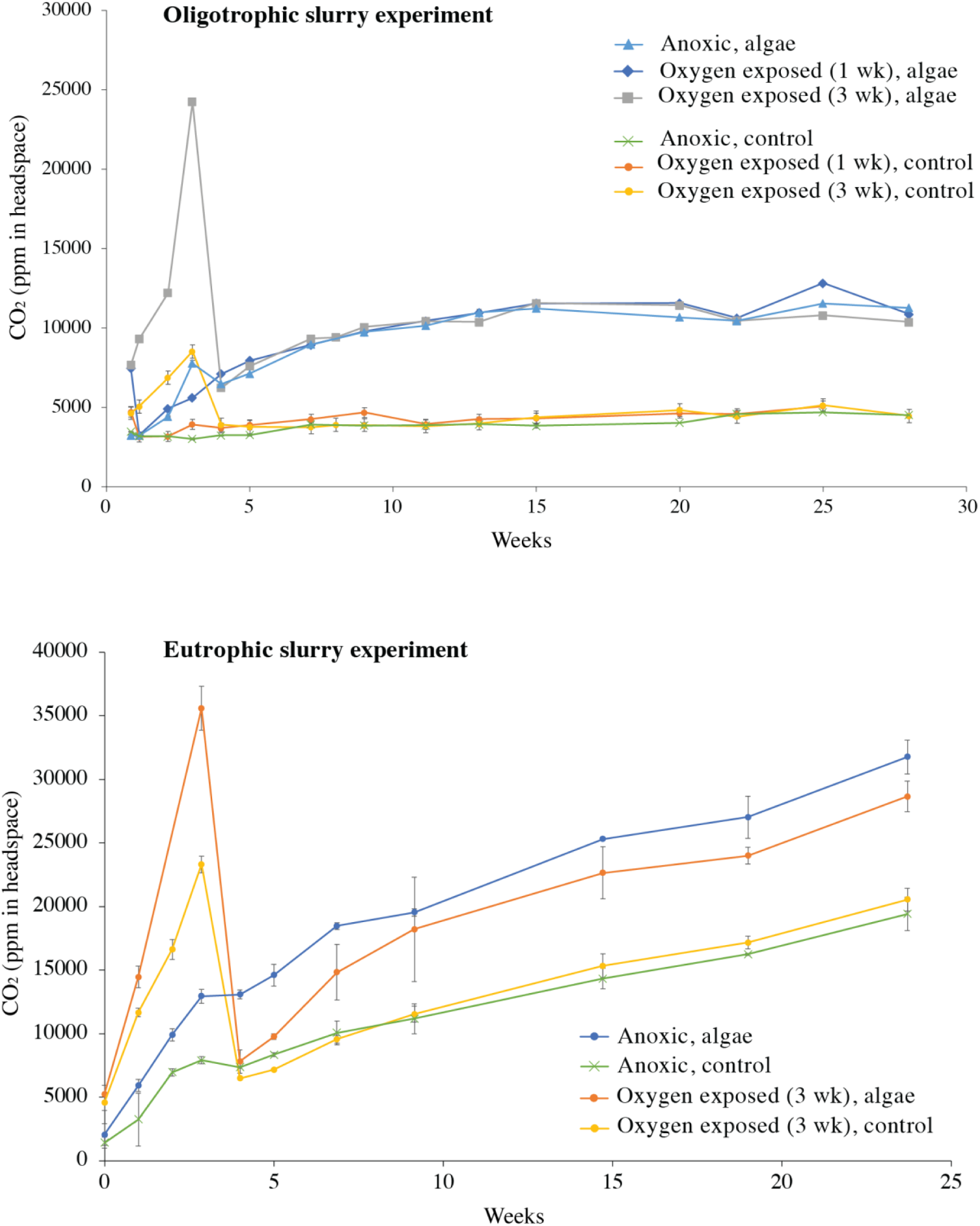
CO_2_ concentrations in the headspace of the slurry experiments. The sudden drop is caused by the flushing of the headspace with N_2_ at the oxic-anoxic switch after 3 weeks.

**Fig. S8.**
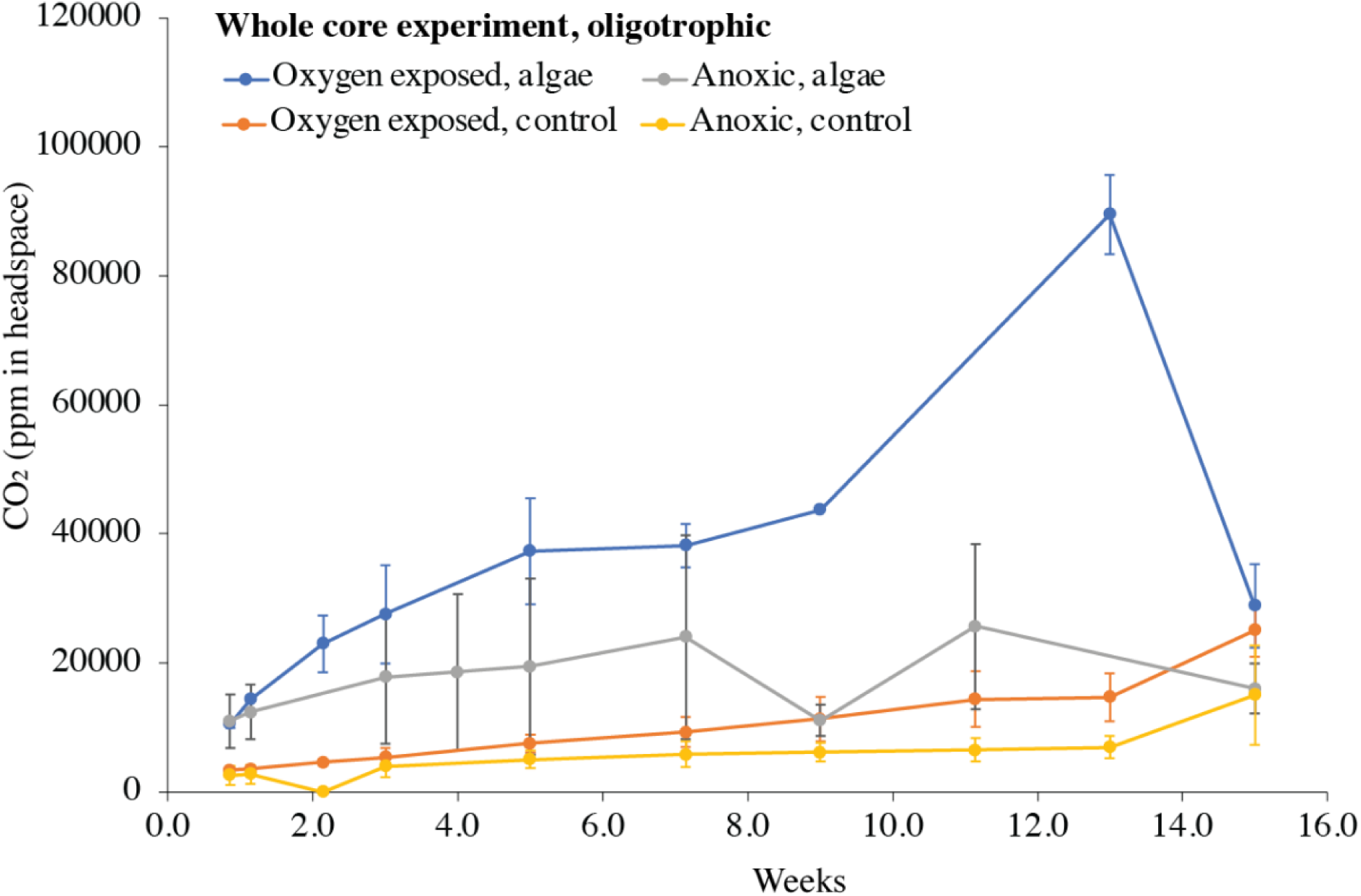
CO_2_ concentrations in the headspace of the whole core experiments.

